# Functional analysis of *Mmd2* and related *PAQR* genes during sex determination in mice

**DOI:** 10.1101/2021.07.10.451886

**Authors:** Liang Zhao, Ella Thomson, Ee Ting Ng, Enya Longmuss, Terje Svingen, Stefan Bagheri-Fam, Alexander Quinn, Vincent R. Harley, Leonard C. Harrison, Emanuele Pelosi, Peter Koopman

**Author notes:** Joint first authors (Liang Zhao, Ella Thomson). Joint last authors (Emanuele Pelosi, Peter Koopman).

## Abstract

Sex determination in eutherian mammals is controlled by the Y-linked gene *Sry*, which drives the formation of testes in male embryos. Despite extensive study, the genetic steps linking *Sry* action and male sex determination remain largely unknown. Here, we focused on *Mmd2*, a gene that encodes a member of the progestin and adipoQ receptor (PAQR) family. We show that *Mmd2* is expressed during the sex-determining period in XY but not XX gonads, specifically in the Sertoli cell lineage which orchestrates early testis development. Analysis of knockout mice deficient in *Sox9* and *Sf1* revealed that *Mmd2* operates downstream of these known sex-determining genes. However, when we used CRISPR to ablate *Mmd2* in the mouse, fetal testis development appeared to progress normally. To determine if other genes might have compensated for the loss of *Mmd2*, we identified the closely related PAQR family members *Paqr8* and *Mmd* as also being expressed during testis development. We used CRISPR to generate mouse strains deficient in *Paqr8* and *Mmd*, but both knockout lines appeared phenotypically normal and fertile. Finally, we generated *Mmd2*;*Mmd* and *Mmd2*;*Paqr8* double-null embryos and again observed normal testis development. These results may reflect functional redundancy among these factors. Our findings highlight the difficulties involved in identifying genes with a functional role in sex determination and gonadal development through expression screening and loss-of-function analyses of individual candidate genes, and may help to explain the paucity of genes in which variations have been found to cause human disorders/differences of sex development.

## Introduction

In most eutherian mammals, male sex determination is regulated by the sex-determining region Y gene *Sry* (Gubbay et al., 1990; Koopman et al., 1991; Sinclair et al., 1990). When expressed in fetal genital ridges, the transcription factor SRY, together with co-factor SF1 (also known as NR5A1), triggers the expression of *Sox9* (Sekido and Lovell-Badge, 2008), which drives the differentiation of Sertoli cells and hence the differentiation of testes (Vidal et al., 2001). In the absence of *Sry* expression or subsequent upregulation of *Sox9*, the bipotential gonads develop instead into ovaries. Despite three decades of research and significant progress in the field, many aspects of the molecular mechanisms regulating fetal testis development remain unclear. As a result, the cause of most 46,XY disorders/differences of sex development (DSD) remains unknown (Eggers et al, 2016).

The progestin and adipoQ receptor (PAQR) family of proteins consists of 11 membrane receptors that are found throughout eukaryotes and in some eubacteria, and show high conservation between human and mouse (Tang et al., 2005). One member of this family, *Mmd2* (also known as *Paqr10*), has previously been identified in a number of screening studies as being expressed in pre-Sertoli cells of the developing testis, but not in the ovary (Menke and Page, 2002; Beverdam and Koopman, 2006; Bouma et al., 2007; Cory et al., 2007; Jameson et al., 2012; Nef et al., 2005; Small et al., 2005; Zhao et al., 2018). In view of these consistent observations, we decided to investigate a possible role for *Mmd2* in testis development.

Previous studies have described roles for *Mmd2* in the developing pancreas and brain. The encoded protein was shown to act in mitochondria to regulate pancreatic endocrine cell development/survival (Gonez et al., 2008). It also functions during gliogenesis, where it was found to localize to the Golgi apparatus, regulating ERK/MAPK signaling by tethering and activating Ras proteins, and maintaining their localization within the Golgi (Jin et al., 2012a, 2012b). The possible link with ERK/MAPK signalling may be relevant in the context of male sex determination, given that a number of regulators of this pathway are known to play a role in testis development (Pearlman et al., 2010; Bogani et al., 2009; Warr et al., 2016). Further, *Mmd2* is regulated by SOX9 to control the proliferation of glial precursors within the spinal cord (Kang et al., 2012), raising the possibility that it may also be regulated by SOX9 during testis development. Accordingly, *Mmd2* is among 1903 genes identified as potential SOX9 targets by chromatin immunoprecipitation of mouse fetal testis extracts (Li et al., 2014), although a direct regulatory relationship was not confirmed experimentally.

In the present study, we studied in detail the expression dynamics of *Mmd2* in the developing mouse fetal gonads and established *Mmd2* as one of the earliest Sertoli cell markers. We confirmed that MMD2 is localized to the Golgi apparatus in Sertoli-like cells and acts downstream of *Sox9*. However, fetal testes appeared normal in mice lacking *Mmd2* alone and in combination with *Mmd* and *Paqr8*; two PAQR family members expressed in the developing gonads. These results may reflect functional redundancy within the PAQR family during gonadal development.

## Materials and methods

### Mice

Mouse embryos were collected from timed matings of the relevant strains, with noon of the day when the mating plug was observed designated 0.5 dpc. Embryos were sexed by PCR (McFarlane et al., 2013) and by morphological assessment of the gonads (12.5–15.5 dpc embryos). To ensure correct staging at 10.5 and 11.5 dpc, tail somites (ts) were counted as described (Hacker et al., 1995): ts 7–9 at 10.5 dpc and ts 17–19 at 11.5 dpc. *W*^*e*^ strain (Buehr et al., 1993), Wt1-RG red-green reporter strain (Zhao et al., 2014b), AMH-Cre;*Sox9*^flox/flox^ mice (Barrionueo et al., 2009), and *Cited2*-knockout strain (Barbera et al., 2002) have been described previously.

To generate the *Mmd2*^flox^ allele, two *lox*P sites were inserted into the *Mmd2* locus flanking exons 3 and 4 by gene targeting (Ozgene). The *Mmd2*^flox^ allele was recombined using a CMV-Cre allele (Schwenk et al., 1995) to generate the *Mmd2*-null allele. The CMV-Cre allele was subsequently excluded from the line by selective breeding.

The *Mmd*-null and *Paqr8*-null alleles were generated using the Alt-R CRISPR/Cas9 system (IDT). Briefly, for each deletion, a pair of crRNAs were separately annealed with tracrRNA. Each duplex RNA was incubated with Cas9 protein to form Cas9 ribonucleoprotein (RNP). Two Cas9 RNP preparations were then mixed 1:1 and subsequently microinjected into B6 one-cell embryos. In the microinjection cocktail, the final concentration of crRNA and tracrRNA was 15 ng/μl with Cas9 protein at 60 ng/μl. Injected embryos were incubated overnight to the two-cell stage and surgically transferred to pseudopregnant CD1 females as described (Zhao et al., 2014a).

Sequences of crRNAs and genotyping primers are described in Supplementary Table 1. All mutant strains were maintained on a C57BL/6 background. Animal experimentation was approved and carried out according to the guidelines established by the University of Queensland Animal Ethics Committee.

### Quantitative RT-PCR (RT-qPCR)

For expression analysis in wild type CD1, AMH-*Cre*;*Sox9*^flox/flox^, or *Cited2*-knockout mice, fetal gonads were collected with mesonephros removed. For expression analysis in *Mmd2*, *Mmd*, or *Paqr8* single or double knock-out gonads, fetal gonads with mesonephroi were pooled for analysis. RT-qPCR analysis was as described previously (Zhao et al., 2015), and primer sequences are provided in Supplementary Table 1.

### *In situ* hybridization (ISH)

Embryos and embryonic gonads were dissected and fixed in 4% paraformaldehyde (PFA) in phosphate-buffered saline (PBS) solution at 4°C overnight and subsequently washed with PBS. Whole-mount ISH was carried out as described (Hargrave et al., 2006). For section ISH, embryos were mounted in paraffin and sectioned at 7 μm using a microtome. Section ISH was carried out as described (Zhao et al., 2017). In situ probe was generated using PCR (primer sequences described in Supplementary Table 1).

### Cell Sorting

Multiple pairs (n >4) of fetal gonads (removed from the mesonephroi) were dissected from Wt1-RG embryos (Zhao et al., 2014b) at 13.5 dpc. Gonadal cells were dissociated and sorted for mCherry and EGFP fluorescence as described (Zhao et al., 2017).

### Immunofluorescence staining of transfected cells

The coding region of mouse *Mmd2* was cloned into pcDNA3 to generate pcDNA-Mmd2. This was transfected into 15P-1 (Paquis-Flucklinger et al., 1993) or TM4 (Mather, 1980) cells, two mouse Sertoli-like cell lines. Transfected cells were immuno-stained as described (Neumann et al., 2008). Primary antibodies were used at the following dilutions: mouse anti-MMD2 (Abcam), 1:100; and rabbit anti-Giantin (Abcam) 1:200.

### Immunofluorescence staining of mouse tissue sections

Immunofluorescence was performed on 7-μm paraffin sections as described (Zhao et al., 2015). Images were captured on a LSM710 confocal microscope (Zeiss). Primary antibodies were used at the following dilutions: goat-anti-AMH (Santa Cruz), 1:500; rabbit anti-HSD3b (Transgenic Inc.), 1:500; goat anti-FOXL2 (NovusBio), 1:100; and mouse anti-SOX9 (Abnova), 1:200.

## Results

### *Mmd2* is specifically up-regulated in Sertoli cells in the developing fetal testis

We first determined the expression of *Mmd2* in male and female mouse gonads from 10.5 to 15.5 dpc by RT-qPCR. *Mmd2* mRNA was undetectable in XX gonads at all developmental stages, whereas XY gonads showed an increasing level of mRNA from 10.5 dpc to 15.5 dpc (Fig 1A). These results were confirmed by whole-mount *in situ* hybridization (WISH) where no expression was evident in the ovaries (Fig 1B). Conversely, *Mmd2* started to be expressed in the undifferentiated testis from around 11.5 dpc, and staining increased in intensity until 15.5 dpc (Fig 1B).

**Figure 1.**
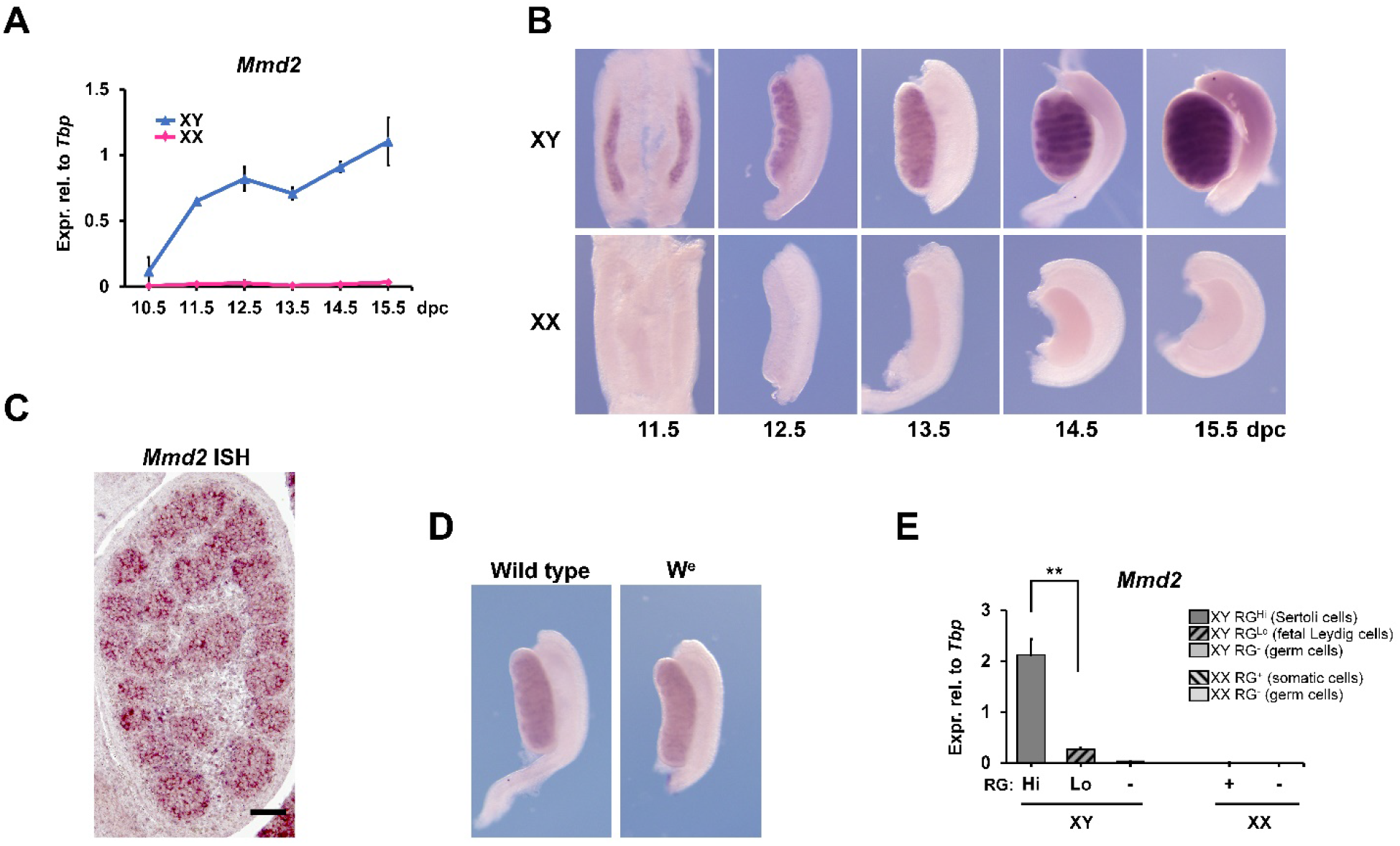
*Mmd2* is specifically up-regulated in Sertoli cells in the developing mouse testis. (A) RT-qPCR showing *Mmd2* expression specifically up-regulated in mouse fetal testis from 11.5 to 15.5 dpc. Mean ± s.e.m., n = 3. (B) Whole-mount in situ hybridisation suggesting *Mmd2* expression restricted in testis cords from 12.5 dpc onwards in fetal testes. n = 3-5 per sex per stage; representative images are shown. (C) Section in situ hybridisation at 13.5 dpc confirming that *Mmd2* expression was predominantly detected within testis cords (n = 3; a representative image is shown). (D) Whole-mount in situ hybridisation showing the presence of *Mmd2* transcripts in the *W*^e^ fetal testis (n = 4; representative images are shown). (E) RT-qPCR analysis on Wt1-RG sorted cell populations confirming that *Mmd2* was predominantly expressed in XY RG^Hi^ cell population enriched for Sertoli cells. Cells from multiple pairs of fetal gonads were pooled and sorted. Mean ± s.e.m. of triplicate RT-qPCR reactions. \*\**P* < 0.01 (Sidak’s multiple comparisons test).

*Mmd2* expression localized to the testis cords (Fig 1B), with *in situ* hybridization signal suggesting that *Mmd2* marks Sertoli cells that are situated between the central germ cells and the peripheral layer of myoid cells (Fig 1C). To address whether *Mmd2* is expressed in germline or somatic cells, we also analysed expression in the *We* mouse strain. This line harbours a mutation in the germ cell marker *c-kit*, resulting in gonads lacking germ cells, but retaining somatic cells. WISH showed no difference in staining between wild type and W*e* XY gonads, confirming *Mmd2* expression within the somatic cells of the testis cords (Fig 1D). To further define the specific cell type expressing *Mmd2*, we used the Wt1-RG red green reporter mouse strain (Zhao et al., 2014b). We previously showed that Sertoli, fetal Leydig and germ cell populations can be identified by sorting via mCherry and EGFP fluorescence, and analysis of specific lineage markers (Zhao et al., 2017). In XY gonads, cells showing strong expression of the RG reporter proteins are Sertoli cells as confirmed by expression of markers including *Sox9* and *Amh*. Cells with low levels of RG are Leydig cells, characterized by the specific expression of *Hsd3b*. RG-negative cells are germ cells, expressing the marker *Mvh* (Zhao et al., 2017). According to these criteria, *Mmd2* transcript was strongly associated with Sertoli cells, with very low levels present in the fetal Leydig population, and none in XY germ cells or any XX gonadal cell populations (Fig 1E). Hence, the timing and cell type-specificity of *Mmd2* expression are consistent with a role in regulating sex determination.

### *Mmd2* expression is dependent on SOX9 and SF1

*Sox9* is one of the major downstream effectors of SRY and plays a critical role in sex determination. To determine whether *Mmd2* acts up- or downstream of SOX9, we used the AMH-*Cre*; Sox9^flox/flox^ mouse line, in which *Sox9* is conditionally deleted in the somatic Sertoli cells. At 13.5 dpc, ablation of *Sox9* led to a 43% reduction of *Mmd2* expression compared to wild type (Fig. 2A), suggesting expression is downstream of, and partially dependent on, SOX9 within these cells. This finding is consistent with previous studies showing that *Mmd2* is a SOX9 target in the spinal cord and testes (Kang et al., 2012; Li et al., 2014).

**Figure 2.**
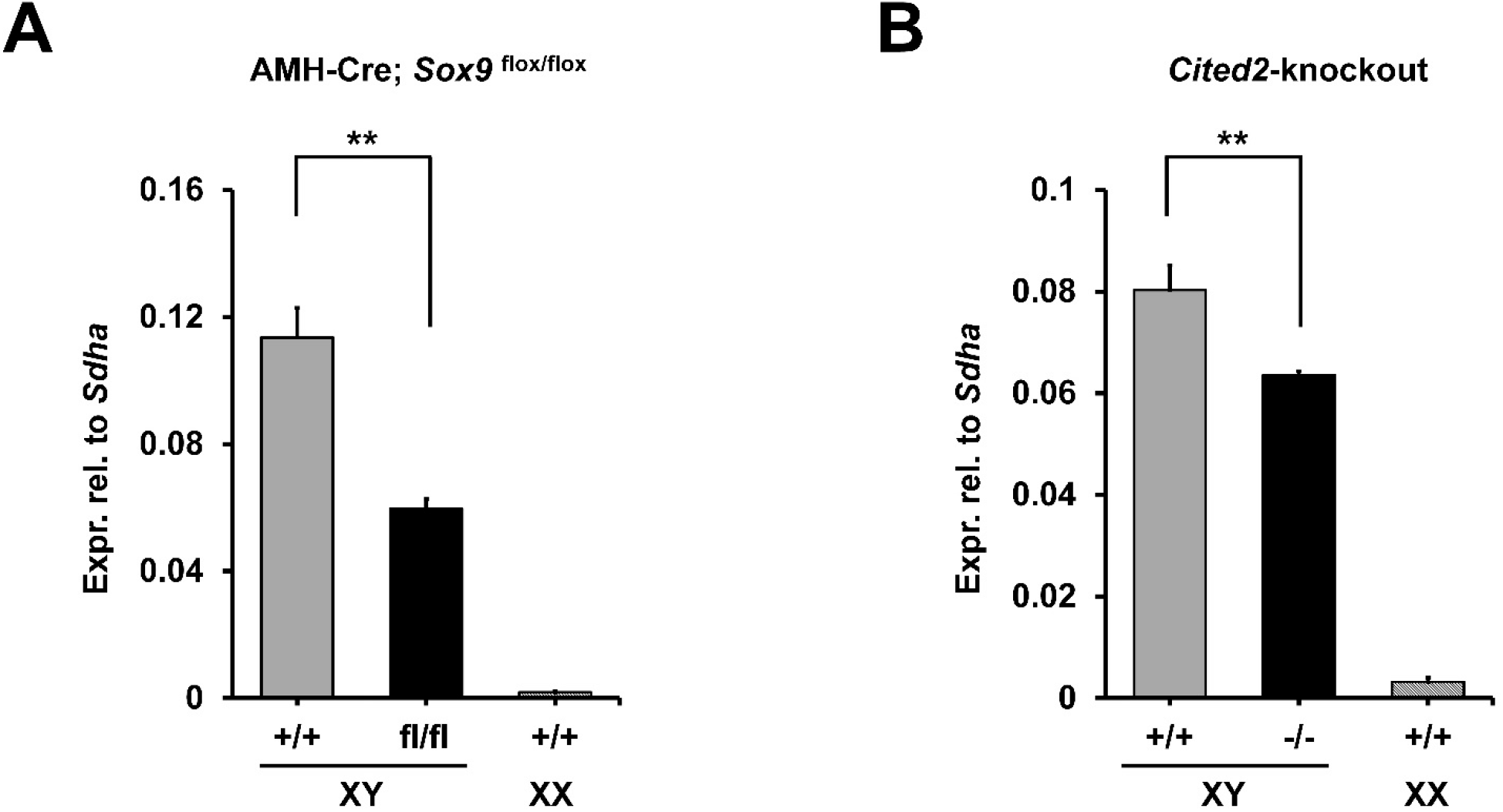
*Mmd2* expression is dependent on *Sox9* and *Sf1*. RT-qPCR analysis showing significant down-regulation of *Mmd2* in either AMH-Cre; *Sox9*^flox/flox^ (13.5 dpc; A) or *Cited2*^−/−^ fetal testes (12.5 dpc; B), which have reduced *Sf1* expression. (A) n = 3 (XY KO), 4 (XY WT), or 5 (XX WT). KO, knockout; WT, wild type. (B) n = 3. Mean ± s.e.m., \*\**P* < 0.01 (Sidak’s multiple comparisons test).

SF1 is an important transcription factor involved in sex determination and development. Together with SRY, SF1 upregulates *Sox9* expression, and acts with SOX9 to activate transcription of a number of genes involved in testis development (De Santa et al., 1998; Wilson et al., 2005; Wilhelm et al., 2007; Sekido et al., 2008; Kashimada et al., 2011). Mice deleted for *Sf1* lack gonads, making them unsuitable for this study (Luo et al., 1994; Sadovsky et al., 1995). Therefore, to determine if *Mmd2* expression was also dependent on SF1, we used the *Cited2*^−/−^ mouse model. Cited2 interacts with the transcription factor WT1 within the embryonic gonads, to regulate the expression of *Sf1* (Buaas and Swain, 2009). A *Cited2−/−* line shows hypomorphic *Sf1* expression that can be used to study the development of the testis (Combes et al., 2010). We found that *Cited2*^−/−^ gonads showed a significant reduction in *Mmd2* expression compared to wild type at 12.5 dpc, suggesting that *Mmd2* is to some extent dependent on, and downstream of, SF1.

### MMD2 localizes to the Golgi complex in Sertoli-like cells

The MMD2 protein has variously been shown to localize to the Golgi complex or to mitochondria (Jin et al., 2012a; Jin et al., 2012b; Gonez et al., 2008). We examined the cellular localization of MMD2 in vitro using the Sertoli-like cell lines 15p-1 and TM4, transfected with an *Mmd2* expression construct. MMD2 co-localized with the Golgi apparatus marker Giantin in both cell lines (Figure 3). However, no co-localization was evident using the mitochondrion-specific dye Mitotracker (data not shown). These results indicate that MMD2 localizes to the Golgi complex of Sertoli cells.

**Figure 3.**
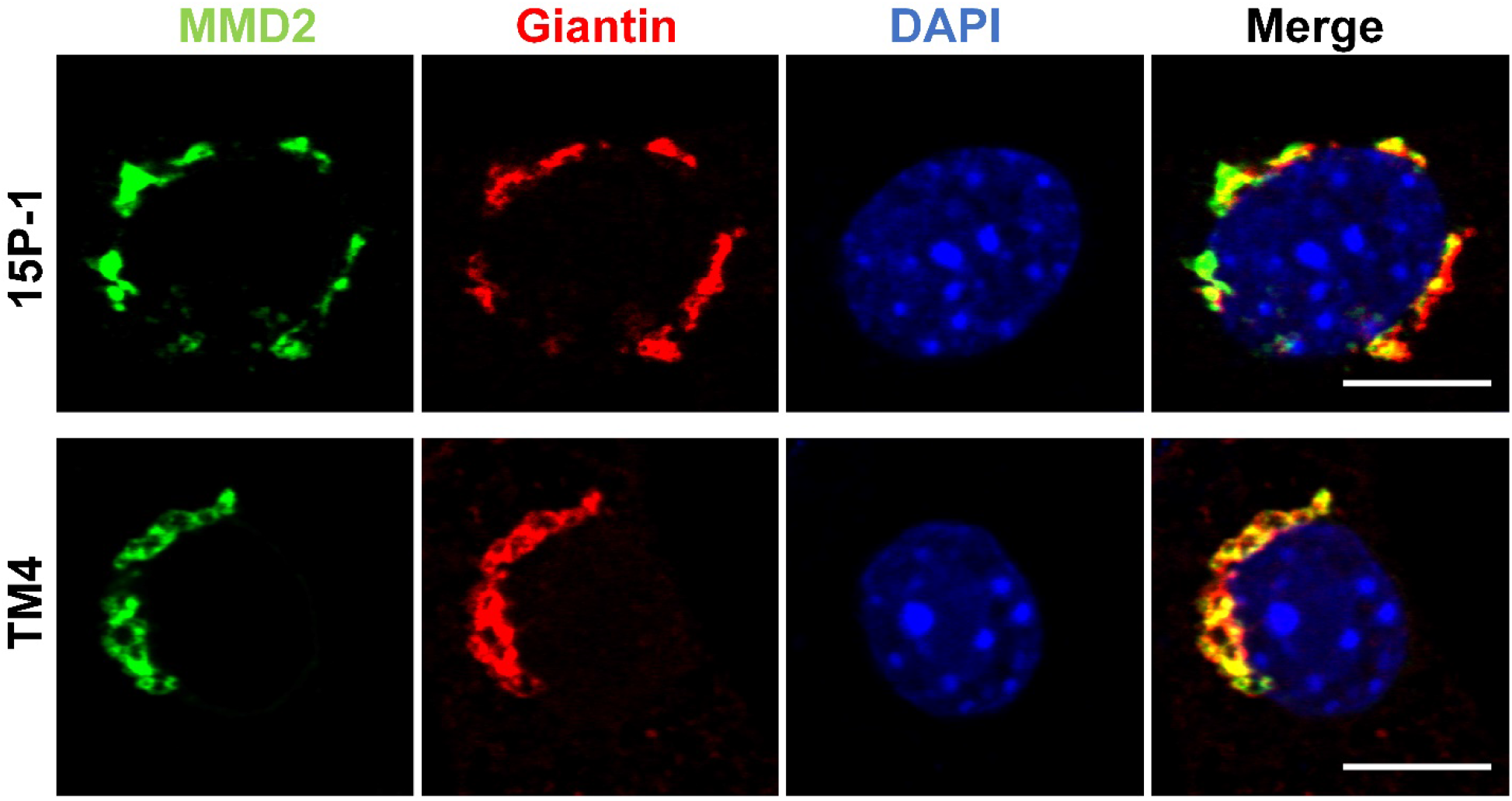
MMD2 localises to Golgi apparatus in mouse Sertoli-like cell lines. MMD2 co-localised with Giantin, a Golgi marker, in transfected 15P-1 or TM4 cells. Scale bar, 10 μm.

### Ablation of *Mmd2* does not affect testis determination and development

To investigate the function of *Mmd2* in testis determination or development, we generated a mouse line in which exons 3 and 4 of *Mmd2* were flanked with loxP recombination sites, and bred these mice with a line ubiquitously expressing Cre recombinase (CMV-Cre; Schwenk et al., 1995) to functionally ablate one (*Mmd2*^+/−^) or both (*Mmd2*^−/−^) alleles (Fig 4A). Gene expression analysis confirmed ablation of *Mmd2*, showing a 50% reduction in intact transcript levels in *Mmd2*^+/−^ XY gonads compared to wild-type, and no detectable expression of intact transcripts in *Mmd2*^−/−^ XY gonads (Fig 4B).

**Figure 4.**
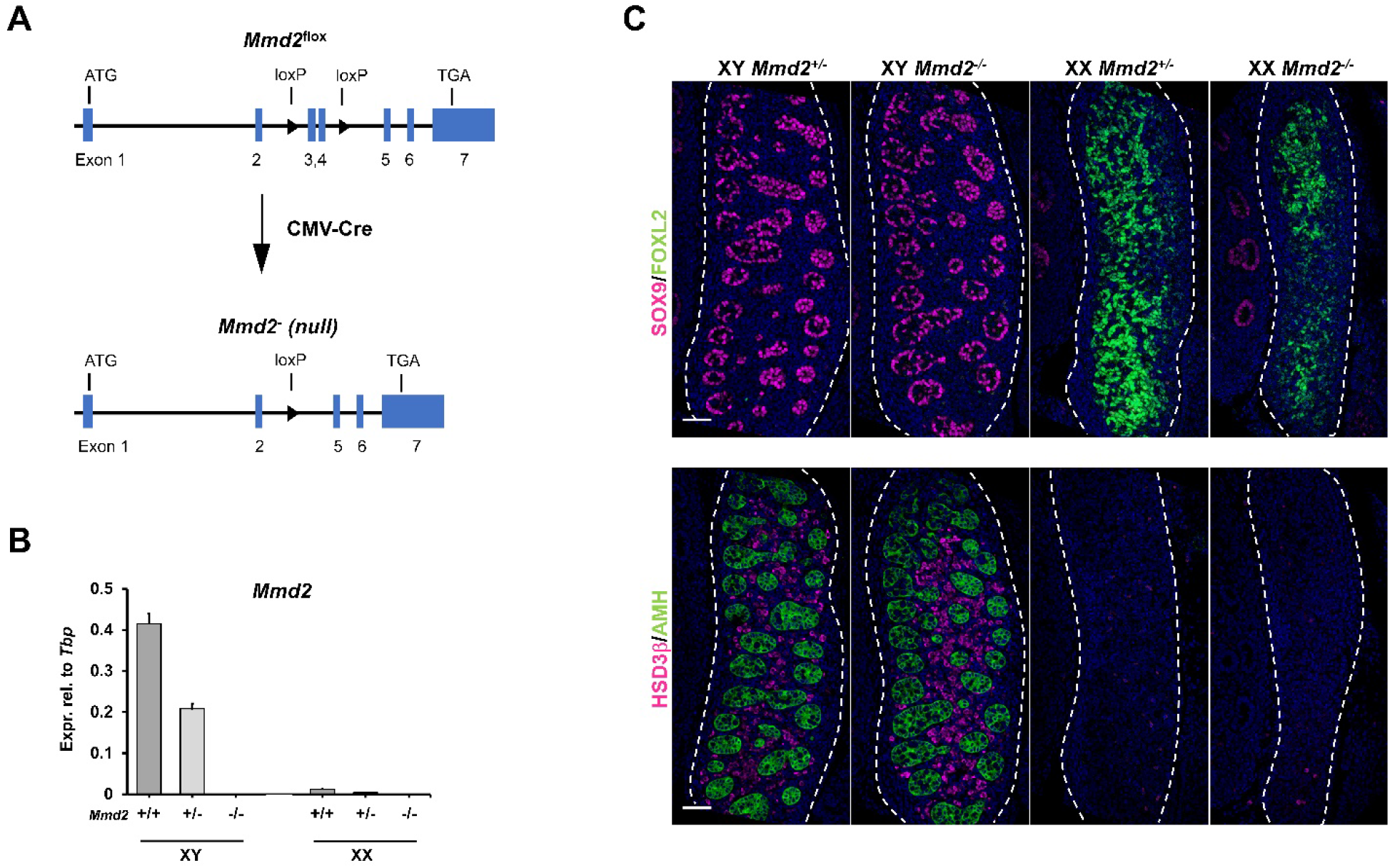
Fetal gonads develop normally in mice lacking *Mmd2*. (A) *Mmd2*^flox^ allele was generated by inserting two loxP sites flanking exons 3 and 4. *Mmd2*^−^ (null) allele was generated by in vivo recombination using the CMV-Cre line. (B) RT-qPCR analysis confirming the absence of *Mmd2* expression in *Mmd2*^−/−^ fetal gonads. Mean ± s.e.m., n = 3 (XX^+/+^; XY^+/−^), 4 (XY^+/+^; XY^−/−^; and XX^−/−^), or 5 (XX^+/−^). (C) Immunofluorescence analysis showing unperturbed expression of sex markers, including SOX9, FOXL2 (upper panel), HSD3β and AMH (bottom panel), in *Mmd2*^−/−^ fetal gonads. *Mmd2*^+/−^ fetal gonads were included as controls. Scale bar, 50 μm.

*Mmd2*^+/−^ and *Mmd2*^−/−^ mice were viable, with no gross phenotypic abnormalities. Embryonic gonads were also grossly normal, with no evidence of gonadal sex reversal or morphological abnormalities. Mutant gonads were collected at 13.5dpc and analyzed by RT-qPCR for expression of a range of lineage marker genes, but no significant differences were seen between wild-type, *Mmd2*^+/−^, and *Mmd2*^−/−^ gonads (Supplementary Figure S1).

Sertoli cell markers SOX9 and AMH were analyzed at the protein level by immunofluorescence. In XY testes, both SOX9 and AMH appeared similar in intensity and localization between *Mmd2*^−/−^ samples and *Mmd2*^+/−^ controls, whereas they were not detected in XX ovaries, as expected (Fig 4C). The Leydig cell marker HSD3B was also assessed and showed no difference in expression between XY *Mmd2*^+/−^ and XY *Mmd2*^−/−^ testes, whereas HSD3B was absent in XX Mmd2+/− ovaries, as expected, and in XX Mmd2−/− ovaries (Fig 4C). Finally, we analyzed the expression of ovarian marker FOXL2 to confirm there was no ectopic development of the granulosa cell lineage within the embryonic *Mmd2*^−/−^ testes. Like in XY *Mmd2*^+/−^ testes, no FOXL2 expression was detected in XY *Mmd2*^−/−^ testes, while FOXL2 was expressed throughout the XX *Mmd2*^+/−^ and *Mmd2*^−/−^ ovaries (Fig 4C). Hence, loss of *Mmd2* caused no detectable impairment of testis development.

### Expression of other PAQR family members in fetal testes

The lack of an obvious phenotype in XY *Mmd2*^−/−^ gonads raised the possibility that other members of the PAQR family might compensate for the loss of MMD2. To test this hypothesis, we focused on two other *Paqr* genes, *Mmd* (*Paqr11*) and *Paqr8*, based on their known function and expression profile. We surveyed our published RNA-Seq dataset derived from analysis of fetal gonads from 10.5-13.5dpc (Zhao et al., 2018), to identify other *Paqr* genes expressed in mouse fetal gonads (Supplementary Figure 2). We found that *Mmd* was robustly expressed in both male and female gonads, peaking between 10.5 and 11.5 dpc, a critical window of sex determination (Supplementary Figure S2). In addition, functional redundancy between MMD2 and MMD has previously been reported in zebrafish and in vitro experiments (Huang et al., 2012; Jin et al., 2012a). A second gene, *Paqr8*, showed consistently higher expression in the male gonad from 11.5 to 13.5dpc, when testis differentiation takes place (Supplementary Figure S2).

We first validated the expression profiles of *Mmd* and *Paqr8* using RT-qPCR on 10.5 to 15.5 dpc gonads. We confirmed that *Mmd* was expressed at similar levels in both XX and XY gonads and was especially strong around the time of sex determination (Fig 5A). Also, in support of RNA-seq data, *Paqr8* was expressed more strongly in XY gonads compared to XX in the period 11.5 – 13.5 dpc (Fig 5B).

**Figure 5.**
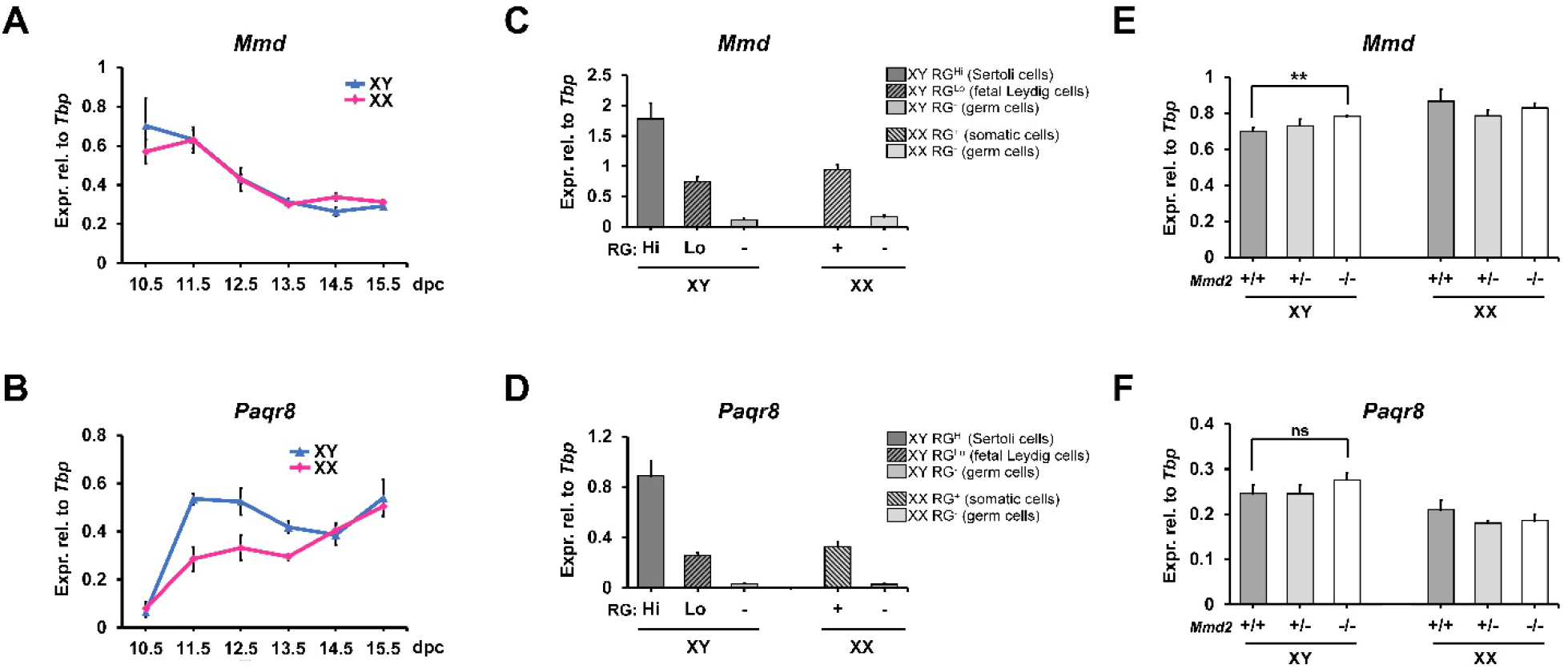
*Mmd* and *Paqr8* are expressed in somatic cells in mouse fetal gonads. (A, B) RT-qPCR showing *Mmd* (A) and *Paqr8* (B) expression up-regulating in mouse fetal testis from 11.5 to 15.5 dpc. Mean ± s.e.m., n = 3. (C, D) RT-qPCR analysis on Wt1-RG sorted cell populations confirming that both *Mmd* and *Paqr8* were expressed in somatic cells in mouse fetal gonads. Within fetal testes, both genes were expressed predominantly in XY RG^Hi^ cell population enriched for Sertoli cells. Cells from multiple pairs of fetal gonads were pooled and sorted. (E, F) RT-qPCR showing expression of *Mmd* (E) and *Paqr8* (F) in gonads from XY and XX mice that were wild type (+/+), heterozygous (+/−) or null (−/−) for *Mmd2*. Mean ± s.e.m. of triplicate RT-qPCR reactions. ** = p <0.01 (Sidak’s multiple comparisons test); ns = not significant.

We then identified the cell types expressing *Mmd* and *Paqr8* using the Wt1-RG mouse line. Like *Mmd2*, both *Mmd* and *Paqr8* were highly expressed in the Sertoli cells of the testis (Fig 5C, D). However, expression was more widespread than *Mmd2*, as both *Mmd* and *Paqr8* were also expressed in fetal Leydig cells and somatic ovarian cells, though at lower levels (Fig 5C, D).

The expression of *Mmd* and *Paqr8* was measured in the *Mmd2* mutant gonads by RT-qPCR to determine if loss of *Mmd2* prompted a compensatory response resulting in their up-regulation. We observed a small but significant increase of *Mmd* expression in *Mmd2*^−/−^ males, but not females (Fig 5E). A trend of increased expression was also observed for *Paqr8* in *Mmd2*^−/−^ testes (Fig 5F), although it was not statistically significant. To investigate if *Mmd* or *Paqr8* are expressed in a SOX9-dependent manner, as is *Mmd2*, we analyzed their expression in the AMH-Cre;*Sox9*^flox/flox^ line. No significant reduction was observed (data not shown), suggesting that *Mmd* and *Paqr8* expression is independent of SOX9 in fetal testes.

### Ablation of either *Mmd* or *Paqr8* function in mice does not affect fetal testis development

To investigate whether *Mmd* or *Paqr8* are required for sex determination or gonadal development, we used CRISPR-Cas9 genome editing to create mouse lines lacking the function of each gene (*Mmd^−/−^*: Fig 6A and *Paqr8^−/−^*: Fig 7A). Gene expression profile analyses using primers specific for the deleted exons confirmed successful disruption of *Mmd* and *Paqr8* respectively (Figs 6B, 7B). Interestingly, we observed an upregulation of *Mmd2* and *Paqr8* in *Mmd*^−/−^ testes, although significant only for *Mmd2* (Fig 6B). Similarly, *Mmd2* and *Mmd* were modestly upregulated in the *Paqr8* testes, but this was not significant (p = 0.054) (Fig 7B).

**Figure 6.**
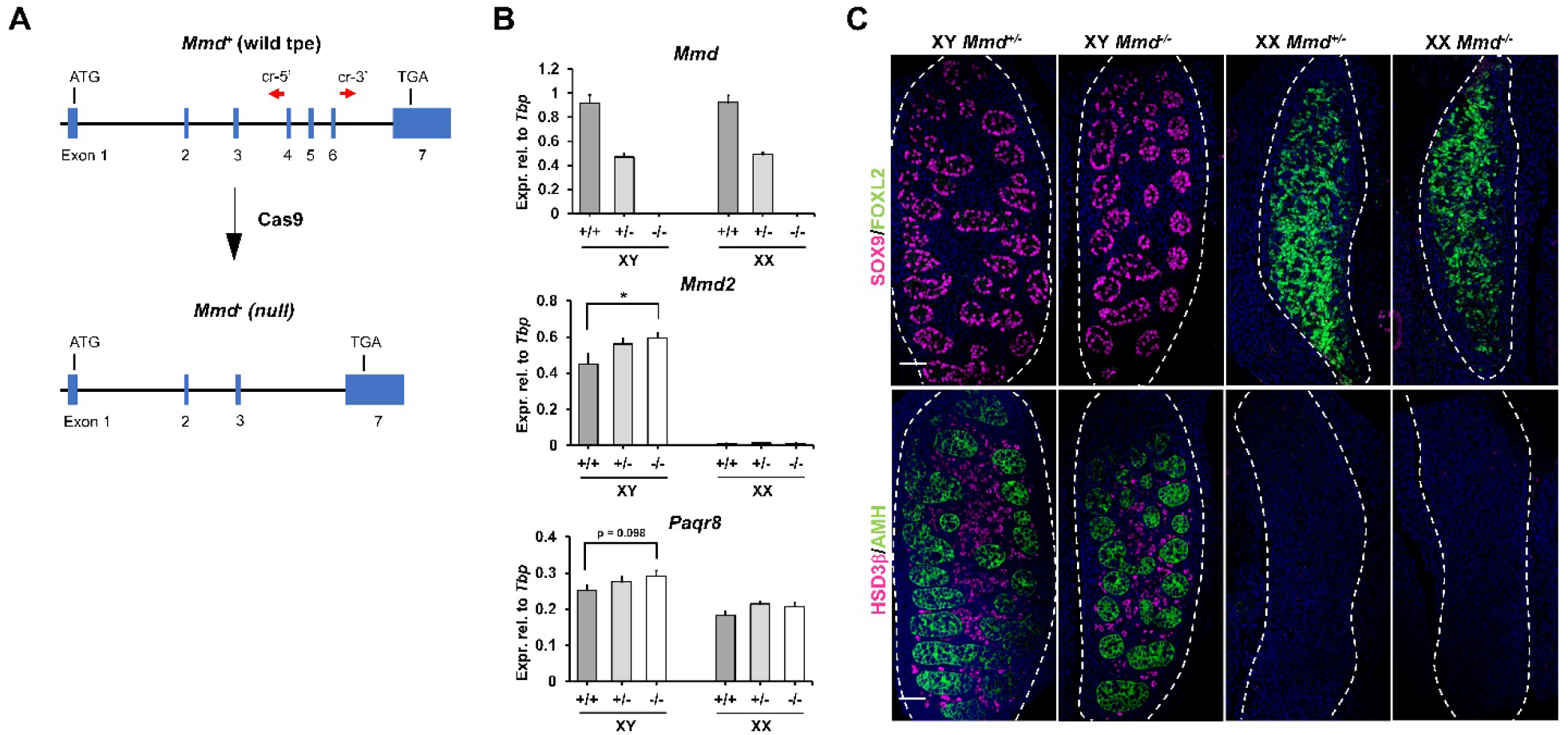
Development of *Mmd*-null fetal gonads appears normal. (A) Generation of *Mmd*^−^ (null) allele via CRISPR in mouse zygotes. cr-5’/3’, crRNA-5’/3’. (B) RT-qPCR analysis confirming lack of *Mmd* expression in the knockout gonads. Note that *Mmd2* and *Paqr8* appeared to be up-regulated in *Mmd*-null gonads. Mean ± s.e.m., n = 3 (XY^+/−^), 4 (XY^+/+^; XX^+/+^; XX^+/−^; and XX^−/−^), or 5 (XY^−/−^). * = p < 0.05 (Sidak’s multiple comparisons test). (C) Immunofluorescence analysis showing similar expression of sex markers, including SOX9, FOXL2 (upper panel), HSD3β and AMH (bottom panel), in *Mmd*^+/−^ and *Mmd*^−/−^ fetal gonads. Scale bar, 50 μm.

**Figure 7.**
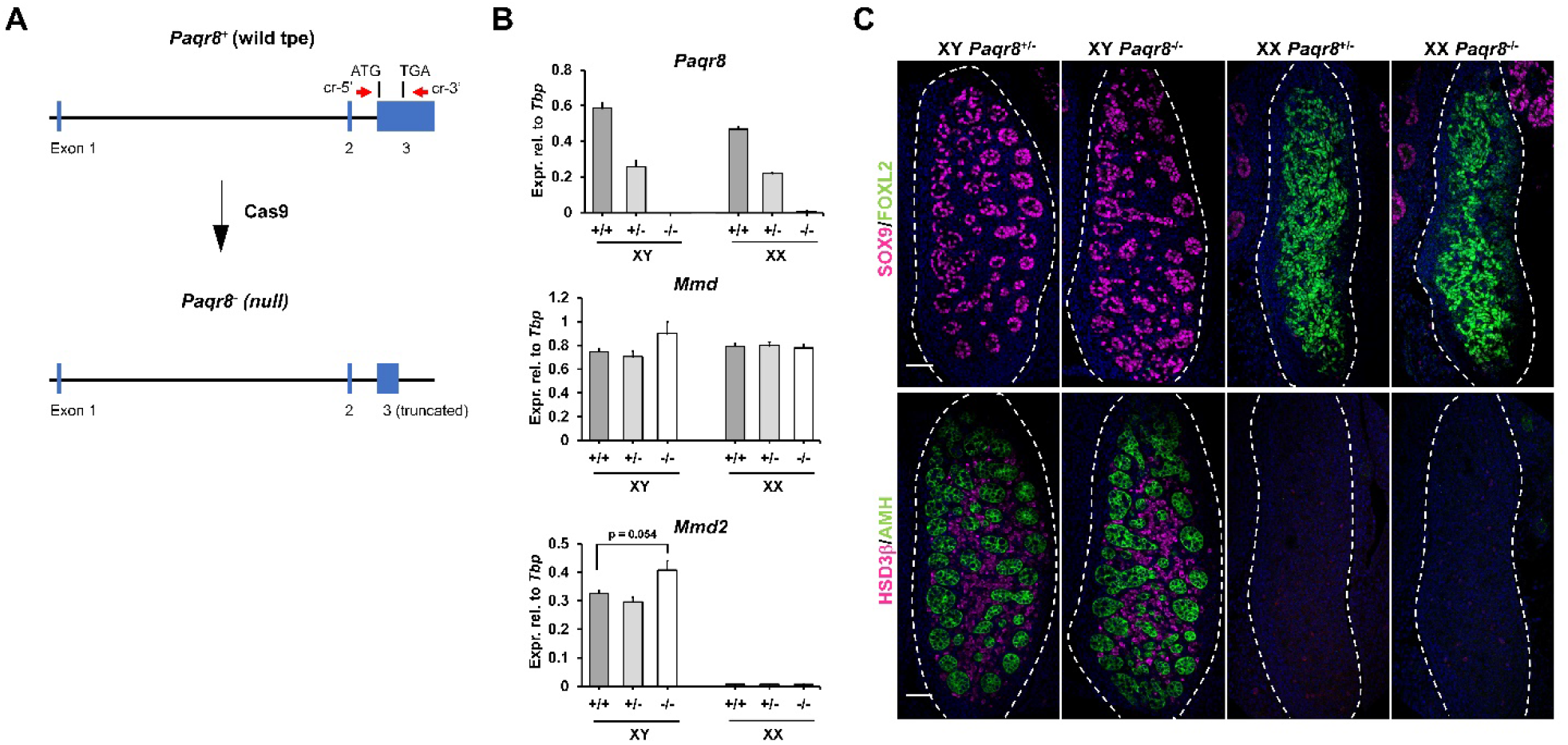
Fetal gonads in *Paqr8*-null mice develop normally. (A) Generation of *Paqr8*^−^ (null) allele via CRISPR in mouse zygotes. cr-5’/3’, crRNA-5’/3’. (B) RT-qPCR analysis confirming lack of *Paqr8* expression in the knockout gonads. Mean ± s.e.m., n = 4. *Mmd* and *Mmd2* were not significantly up-regulated in *Paqr8*-null fetal testes. (C) Immunofluorescence analysis showing similar expression of sex markers, including SOX9, FOXL2 (upper panel), AMH, and HSD3β (bottom panel), in *Paqr8*^+/−^ and *Paqr8*^−/−^ fetal gonads. Scale bar, 50 μm.

Loss of *Mmd* or *Paqr8* function did not cause any gross defects in gonad development in either sex. Expression of a wide range of marker genes was unaffected, as assessed by RT-qPCR (Supplementary Figures S3, S4). Immunofluorescence analysis of key gonadal differentiation markers did not reveal any significant difference between *Mmd*^−/−^ or *Paqr8*^−/−^ gonads and controls (Figs 6C, 7C). We conclude that each of these factors individually is, like *Mmd2*, dispensable for sex determination and gonadal development.

### Combined deletion of *Mmd2* and *Mmd* or *Paqr8* function does not disrupt fetal testis development

Although mice lacking the function of either *Mmd2*, *Mmd*, or *Paqr8* showed no gonadal defects, it remained possible that combined loss-of-function of these genes might affect sex development. To address this possibility, we generated *Mmd2*;*Mmd* and *Mmd2*;*Paqr8* double knockout embryos by breeding the relevant mouse lines, and analyzed their gonads at 13.5dpc. Gene expression analysis confirmed the expected genetic disruption in each case (Fig 8A).

**Figure 8.**
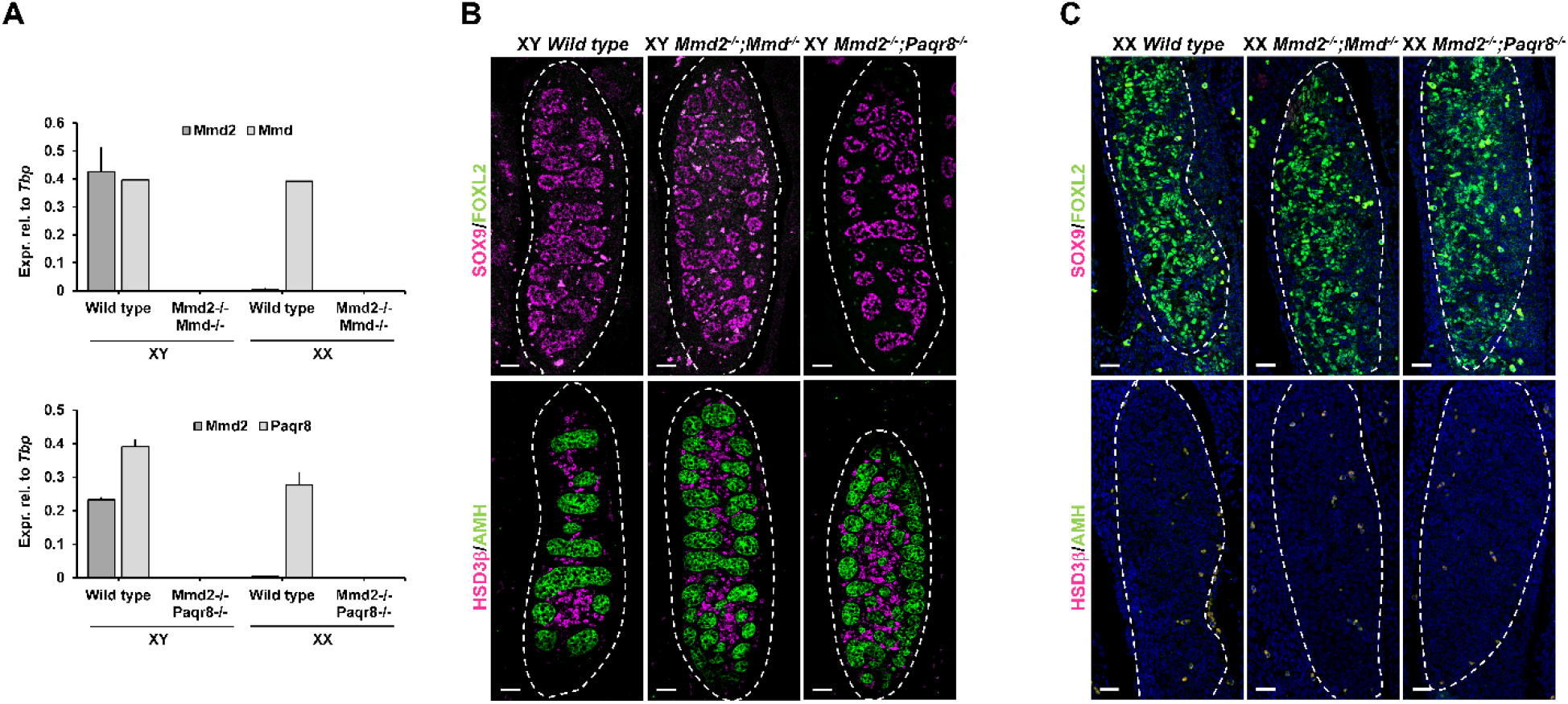
Double knockout of *Mmd2*/*Mmd* and *Mmd2*/*Paqr8* does not disrupt sex determination. (A) RT-qPCR analysis confirming ablation of *Mmd2* in combination with *Mmd* or *Paqr8* in double knockout gonads. Mean ± s.e.m., n = 4. (B, C) Immunofluorescence analysis showing similar expression of sex markers, including SOX9, FOXL2 (upper panel), HSD3β and AMH (bottom panel), in *Mmd2*/*Mmd* and *Mmd2*/*Paqr8* double knockout XY (B) and XX (C) gonads compared to wild type controls. Scale bar, 50 μm.

However, similar to the three singly-targeted lines, no gross fetal gonadal phenotype was observed. Immunofluorescence analysis was performed to assess the expression of the Sertoli cell marker SOX9 and the granulosa cell marker FOXL2 in 13.5 dpc gonads (Fig 8B). No difference was observed between double homozygous deleted mice and double heterozygous controls, and this was further confirmed by analysis of the expression profile of a range of male and female markers including *Sox9*, *Hsd3b*, *Amh*, *Cyp11a1*, *Foxl2*, and *Fst* (Supplementary Figures S5, S6). These results indicate that the combined ablation of *Mmd2* with either *Mmd* or *Paqr8* does not affect sex determination or early gonadal development in the mouse.

## Discussion

Despite extensive study since the discovery of *Sry*, our understanding of the regulatory networks of sex determination in mammals remains patchy, with relatively few functionally important genes having been identified. As a result, many cases of human disorders/differences of sex development (DSD) remain undiagnosed. Efforts to fill in the blanks and assemble genes into pathways have mostly revolved around molecular strategies aimed at identifying targets of key transcription factors known to play a role in gonadal development (Li et al., 2014, Garcia-Ortiz et al., 2009), genomic analysis of DNA from individuals with DSD to identify causative variants (Croft et al., 2018, Sreenivasan et al., 2018), and expression screening strategies in model organisms such as mice (Jameson et al., 2012, Zhao et al., 2018).

The logic underpinning expression screening appears robust and is summarized as follows. If a gene is expressed in the developing testes but not ovaries, or vice versa, and/or if its expression is confined to a time interval in which critical sex determination or differentiation decisions are occurring, then such a gene is likely to play a functional role in those events. This likelihood is further increased if the gene can be shown to associate with a cell lineage known to orchestrate key events in sex differentiation, in this case the Sertoli cell lineage.

In this study, we focused our attention on *Mmd2*, a gene that we found to match all of these criteria. It was therefore surprising to find that loss of *Mmd2* function had no discernible effect on sex determination or the development of testes in mice. Although we cannot exclude the possibility that *Mmd2* expression is functionally unrelated to early testis development, the logic described above led us to seek other explanations for the lack of phenotype. We reasoned that lack of MMD2 may be compensated within the testes by the expression of other PAQR family members. To test this hypothesis, we generated null mouse lines for the related genes *Mmd* and *Paqr8* using CRISPR. *Mmd* is functionally redundant with *Mmd2* in the developing heart in zebrafish (Huang et al., 2012), and *Paqr8* was chosen due to its expression profile showing a greater upregulation in XY gonads at the time of sex determination – a profile unique among the *Paqr* genes. Both genes were co-expressed with *Mmd2* during testis development. Neither *Mmd*^−/−^ nor *Paqr8*^−/−^ mice showed evidence of sex reversal or disruption of gonadal development, and double ablation of *Mmd2*/*Mmd* and *Mmd2*/*Paqr8* did not result in impaired sex development. Therefore, deletion of the *Paqr* candidates most likely to play a role in sex determination and gonadal differentiation, either singly or pairwise with *Mmd2*, had no discernible effect in this system.

It is possible that some impairment or phenotypic changes occurred that we were unable to detect in the mutant mice. For each strain generated in this study, we applied a comprehensive battery of qualitative and quantitative markers of gonadal development. However, gonadal development is strongly canalized, and so it remains formally possible that the program of gonadal development was weakened, but perhaps only transiently and not so much as to cause a failure to meet a threshold required for sex reversal or gonadal dysgenesis. The loss-of-function mutants were created on a C57BL/6 genetic background, which is known to be susceptible to perturbations of the sex determination program (Correa et al., 2012), suggesting that this possibility is unlikely. Nonetheless, it remains possible that an overt phenotype might have been exposed by crossing the mice with other partially compromised strains such as *Sox9* heterozygous mutant mice (Barrionuevo et al., 2006; Bagheri-Fam et al., 2008).

We consider it most likely that functional redundancy exists between the three factors. While *Mmd2* and *Mmd* have been shown to act redundantly in the zebrafish heart (Huang et al., 2012), loss of both together did not compromise mouse gonadal development. This issue may be resolved by the creation of triple mutant mice deficient in *Mmd2*, *Mmd* and *Paqr8* together. Even so, it remains possible that other PAQR factors might be capable of masking a phenotype in the triple mutants. Our analysis showed that several *Paqr* genes are expressed during gonadal development in mice, notably *Paqr7,* which shows some degree of preferential expression in testes, and *Paqr1* and −*2*, which are expressed at a moderate level in gonads of both sexes (Supplementary Figure S2; Zhao et al., 2018). In addition, it is possible that one or more *Paqr* genes would show compensatory upregulation in the absence of MMD2, MMD, and PAQR8 or combinations thereof, consistent with the compensatory upregulation among *Mmd*, *Mmd2* and *Paqr8* detected in the present study. Certainly, our findings highlight the difficulties involved in identifying members of multi-gene families with a functional role in sex determination and gonadal development through loss-of-function analyses of individual genes.

The localization of the MMD2 protein to the Golgi apparatus within the Sertoli cells may reflect a role in regulating the MAPK signaling pathway within the gonad, consistent with the known role of the Golgi as a regulatory node for various signaling cascades (Wei and Seeman, 2009). Further, MMD2, MMD and PAQR8 have previously been found to be involved in MAPK activation by regulating MAPK1/3 phosphorylation (Liu et al., 2012; Kasubuchi et al., 2017). MAPK signaling is critically important for the proper expression of the male-determining gene, *Sry,* which is activated at 10.5 dpc within the pre-Sertoli cells. Targeted loss of components within this pathway, including MAP3K4, MAP2K, and GADD45γ, have resulted in various levels of dysregulation of *Sry*, producing impaired sex development ranging from ovotestis formation to complete XY sex reversal (Bogani D 2009; Warr et al., 2016; Warr et al., 2012). If MMD2, MMD, and PAQR8 do act redundantly in this system, this may reflect regulation of a critical pathway that needs to be protected from the effects of single gene mutation.

Consistent with this concept, we note that a recent study investigated the role of 30 genes that were highly enriched in the testis and believed to be important for testis development and fertility (Lu et al., 2019). The authors found no obvious phenotype nor fertility defects when single knock-out mice were generated, suggesting a high degree of functional redundancy in this system.

The results of our study may also help to explain the paucity of DSD genes that have been positively identified by variant analysis. Currently, around 60% of 46,XY DSD cases have an unknown genetic cause (Eggers et al., 2016), and the identity and role of relevant genes and pathways are still poorly understood. As our understanding of complex disorders increases, it is becoming clear that many congenital conditions have a multigenic component. Regulatory networks with a high degree of redundancy are likely to mask potentially deleterious mutations. Further, generation of mouse models can be complicated by the known differences in dosage sensitivity to loss-of-function thresholds between mice and humans (Barrionuevo et al., 2006, Uda et al., 2004). More sensitive “omics” methods, including improved computational strategies, the ability to generate more genetically complex mouse models using CRISPR technologies, and/or the development of appropriate high-throughput cell-based assays may hold the key to illuminating the spectrum of molecular causes of human DSD.

## Acknowledgements

We thank Dr Johnny Huang (Queensland Facility for Advanced Genome Editing) for zygote microinjections. Confocal microscopy was performed at the Australian Cancer Research Foundation/Institute for Molecular Bioscience Cancer Biology Imaging Facility. This work was supported by grants from the Australian Research Council and the National Health and Medical Research Council (NHMRC) of Australia. PK received salary support as a Senior Principal Research Fellow of the NHMRC.

