## Supplemental Table 1 for "Functional analysis of *Mmd2* and related *PAQR* genes during sex determination in mice"

**Supplementary Table 1**
Oligonucleotide sequences used in experiments.

### ISH probe generation

| Primer | Primer sequence |
|---|---|
| ishMmd2.F5 | CGCGGCGATGTTCACTCTGG |
| ishMmd2.R5 | ACACCCTTGCCCACACCCATCT |

### CRISPR

| crRNA | crRNA sequence |
|---|---|
| crMmd-5' | CATAACCGGACGTGAGATGGGGG |
| crMmd-3' | AGTTACACTAGCTATCCCCAAGG |
| crPaqr8-5' | gGTTGTCTAACTGAGCCCCGAGG |
| crPaqr8-3 | CAGACAGGGGACCGTGGCGCTGG |

### Genotyping PCR

| Primer | Primer sequence |
|---|---|
| genoMmd2.F | GTGTGTTTCCCAGCACCTTTGTCT |
| genoMmd2.R | GACGAGCGTGTTTTCTTACCTACT |
| genoMmd.F | GGGCAAGCAGGAGCTGGTAAGAGAT |
| genoMmd.R | TGGGCCACCACAACCTTCAAAGAT |
| genoMmd.QF1 | CCCCACCCCCAGTTTGGTTTAC |
| genoMmd.QR1 | AGGGGGAAGGGAAGCCTGCTA |
| genoPaqr8.F | CCGTGCTTCCTGGGGTTATCTGAG |
| genoPaqr8.R | GCGGAGGGTGGCTGTTCTGTTAG |
| genoPaqr8.QF1 | AGCGCCCTGGCTCACTTCTTCTA |
| genoPaqr8.QR1 | TAAGGCCGTCGGTAGCGATACTTG |

### qRT-PCR

| Primer | Primer sequence |
|---|---|
| qMmd2.F | GAGTCCCGGCGCACAAGAGG |
| qMmd2.R | CTGCCAAGGATGCTGGGAATGAT |
| qMmd_F | GAACTTTTCTTCTATCTCACGATGG |
| qMmd_R | CTGAAGTCCGTCAGTGTTATTCA |
| qPaqr8.F | TGGAACACTACAGACGCTAATCTC |
| qPaqr8.R | CTCTAGGATGGCAGTCGTCAT |
