## Supplemental Figures for "Functional analysis of *Mmd2* and related *PAQR* genes during sex determination in mice"

### Supplementary Figures

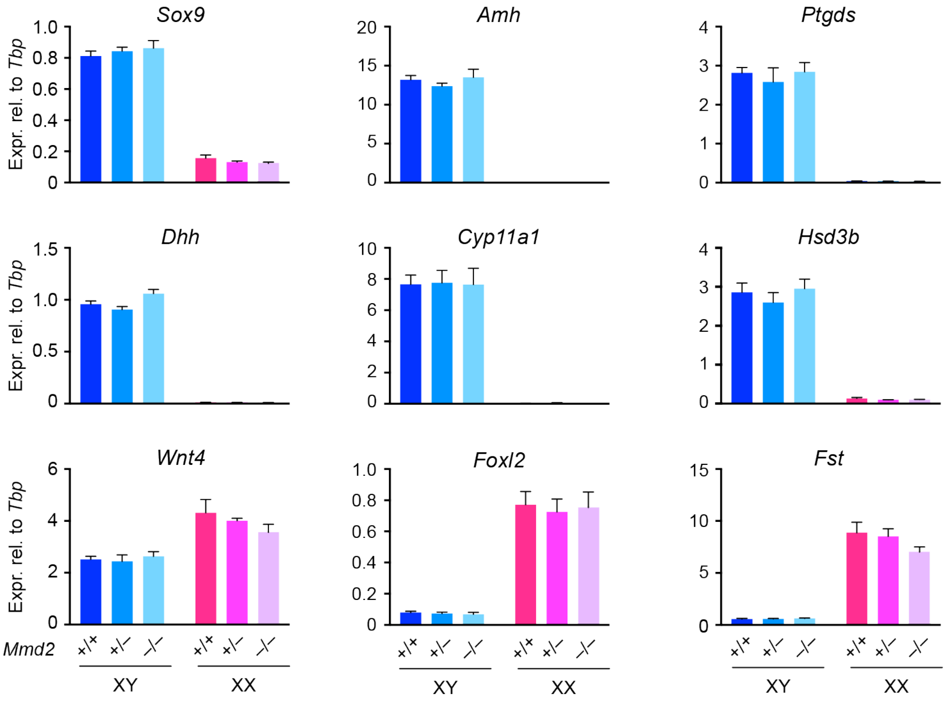


**Supplementary Figure S1.** Unaltered expression of a range of marker genes in *Mmd2*-null fetal gonads. RT-qPCR analysis was performed on *Mmd2* wild type (+/+), heterozygous (+/–) or homozygous (–/–) fetal gonads. Mean ± s.e.m., n = 3 ($XX^{+/+}$; $XY^{+/-}$), 4 ($XY^{+/+}$; $XY^{-/-}$; $XX^{-/-}$), or 5 ($XX^{+/-}$).

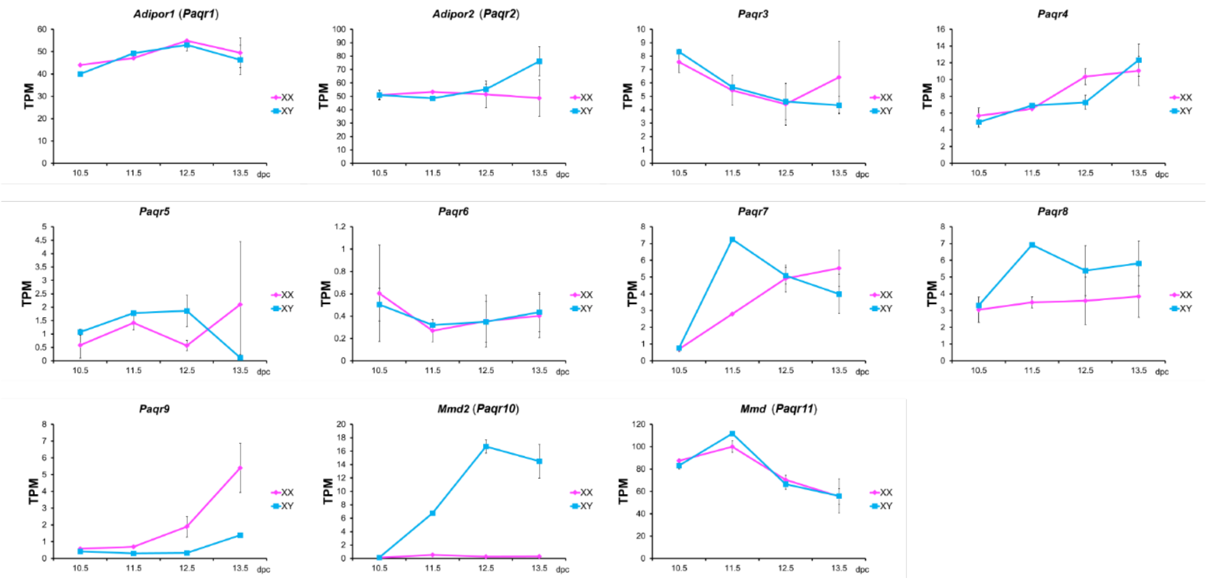


**Supplementary Figure S2.** RNA-Seq expression profiles of Paqr family genes in the developing mouse fetal gonads from 10.5 to 13.5 dpc. Data adapted from our published RNA-Seq data set (Zhao et al., 2018).

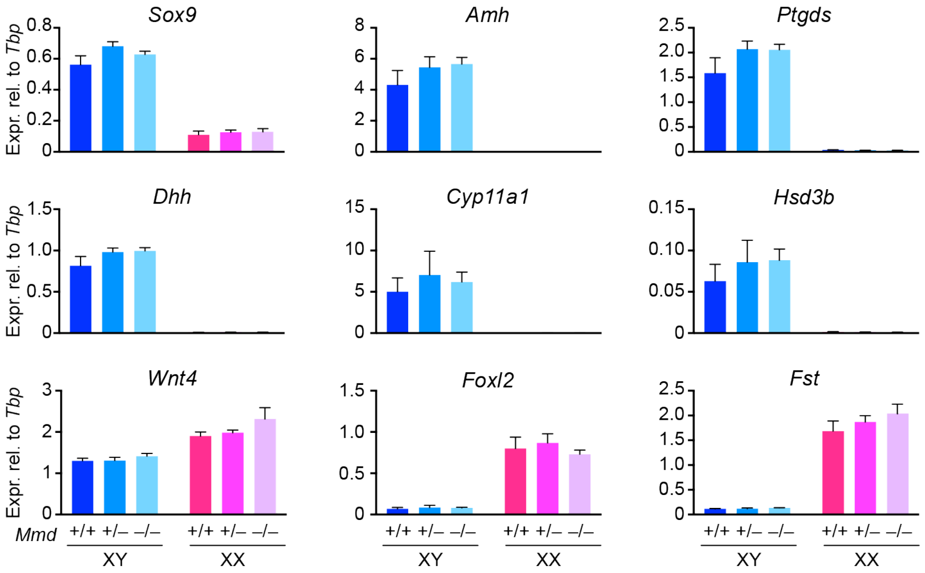


**Supplementary Figure S3.** Unaltered expression of a range of marker genes in *Mmd*-null fetal gonads. RT-qPCR analysis was performed on *Mmd* wild type (+/+), heterozygous (+/–) or homozygous (–/–) fetal gonads. Mean ± s.e.m., n = 3 ($XY^{+/–}$), 4 ($XY^{+/+}$; $XX^{+/+}$; $XX^{+/–}$; and $XX^{–/–}$), or 5 ($XY^{–/–}$).

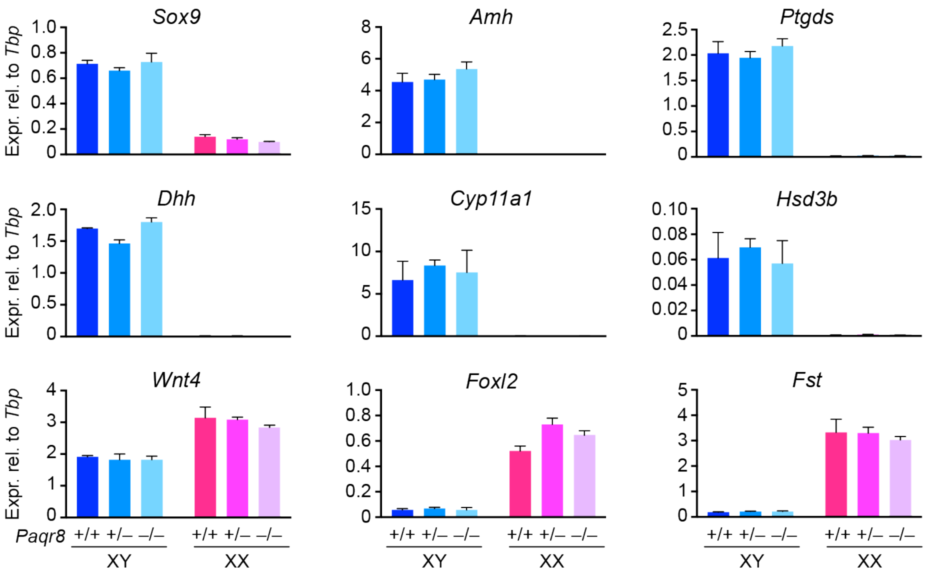


**Supplementary Figure S4.** Unaltered expression of a range of marker genes in *Paqr8*-null fetal gonads. RT-qPCR analysis was performed on *Paqr8* wild type (+/+), heterozygous (+/–) or homozygous (–/–) fetal gonads. Mean ± s.e.m., n = 4.

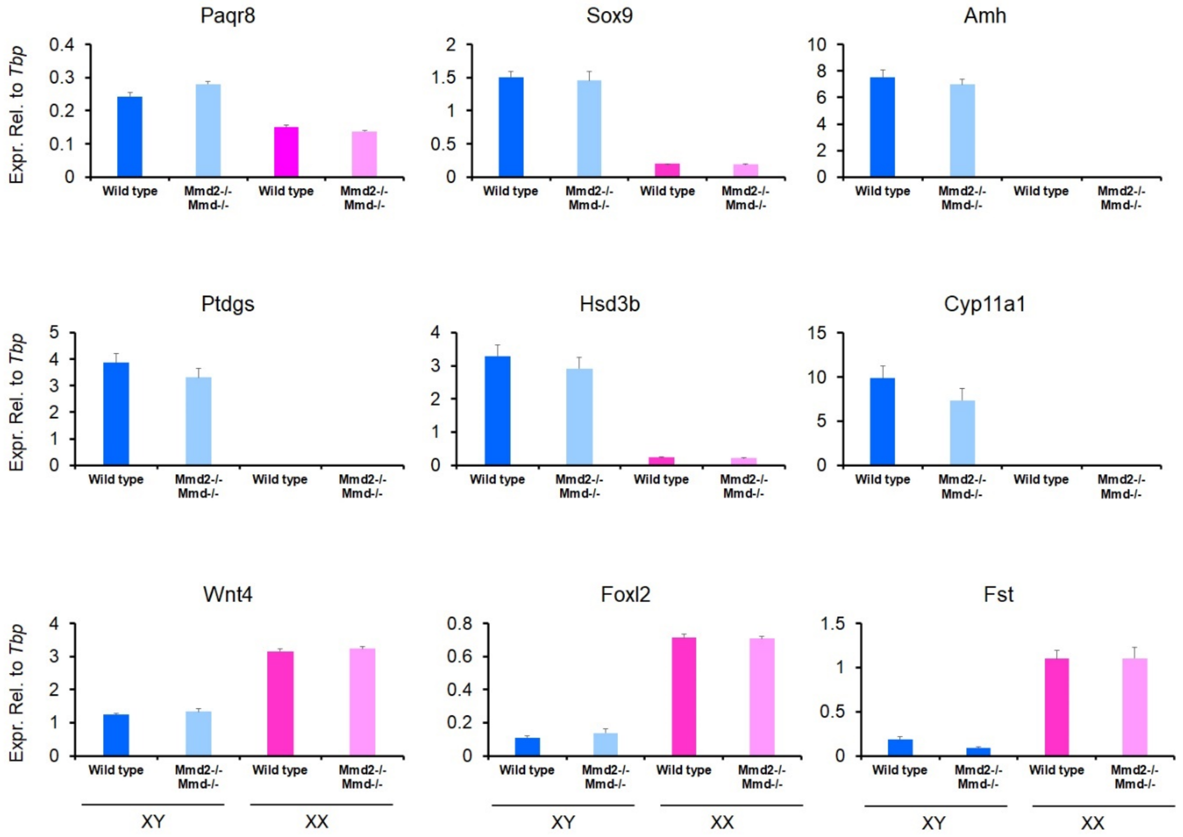


**Supplementary Figure S5.** Unaltered expression of selected marker genes in *Mmd2*/*Mmd* double knockout (*Mmd2*$^{-/-}$;*Mmd*$^{-/-}$) fetal gonads compared to wild type controls. Mean ± s.e.m., n = 4.

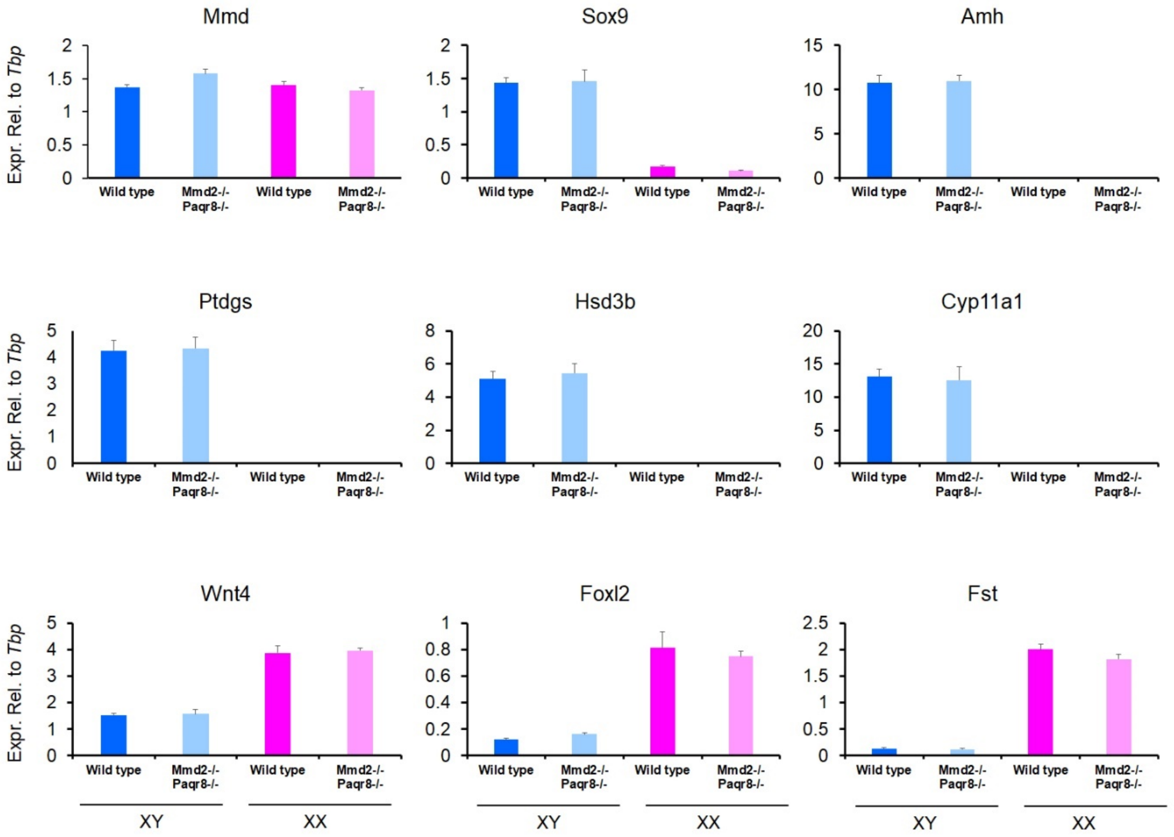


**Supplementary Figure S6.** Unaltered expression of selected marker genes in *Mmd2*/*Paqr8* double knockout (*Mmd2*$^{-/-}$;*Paqr8*$^{-/-}$) fetal gonads compared to wild type controls. Mean ± s.e.m., n = 4.
